# Roles of K-channel activity in feather bud morphogenesis

**DOI:** 10.1101/2025.09.28.679061

**Authors:** Madison Zitting, Zhou Yu, Ting-Xin Jiang, Ping Wu, Randall Widelitz, Cheng-Ming Chuong, Robert Hsiu-Ping Chow

## Abstract

Bioelectricity plays a key role in shaping tissues during early development. In developing chick skin, we previously demonstrated that a transient standing electrical current loop forms within individual feather buds as they start to elongate. Current flows into the feather bud tip via calcium channels, facilitating collective distal dermal cell movement, thus determining bud orientations. Here, we further evaluate the hypothesis that potassium channels carry the current that flows out of the bud base. We found potassium channel blockers convert periodic feather primordia into horizontal stripes and alter bud aspect ratios by disrupting the bud elongation process. Bioelectric characterization showed the entire current loop is altered, not only the outward current at the base, but also the inward current at the feather bud tip. In situ hybridization shows beta-catenin and Sonic Hedgehog, both feather bud markers, are reduced, resulting in altered periodic patterning and bud shaping. These findings show that a complex multiscale interdependency of ion channel function across a multicellular cell collective is essential for periodic feather patterning and shaping of 3-dimensional feather architecture during skin morphogenesis.

**Summary Statement:** Calcium channels at the feather bud tip and potassium channels at the base maintain a standing current loop, creating an essential multicell collective for proper feather bud patterning and morphology.

## Introduction

Tissue patterning is fundamental to tissue development and regeneration (Ramos et al., 2024). We have been using avian embryo skin explants as a model system to study the development and morphogenesis of skin and skin appendages. The advantages are the distinct geometric feather-bud arrangement, well-defined shape of each feather bud, clearly characterized developmental stages (Chuong et al., 2025; Jiang et al., 2004), and the ease of measurements and manipulation (Jiang et al., 2023). In the developing dorsal chicken skin, epithelial-mesenchymal interactions lead to the periodic emergence of feather primordia the morphogenetic field around Hamburger & Hamilton stage 31 (HH31) (Hamburger and Hamilton, 1951), starting at the midline and expanding bilaterally (Chuong et al., 2025; Curantz et al., 2022; Dhouailly, 2024; Ho et al., 2019; Inaba et al., 2019; Turing, 1952). Each feather primordium, consisting of an epithelial placode and subjacent dermal condensation, emerges progressively (Inaba et al., 2019; Riddell and Headon, 2025). Bud dermal cells form a functional syncytium through gap junction-coupling. The short feather bud elongates into a long bud, establishing anterior-posterior and proximal-distal axes (Li et al., 2017; Yu et al., 2002). Although our previous study showed calcium activity is related to collective cell movement, the mechanisms underlying the formation of the bioelectric current loop and the specific channels involved remain unexplored.

This study investigates the function of potassium channels in skin morphogenesis, focusing on two key stages: the early periodic patterning of dorsal skin bud populations and the later shaping of individual feather buds. The importance of biochemical signaling through the Turing reaction-diffusion mechanism uses secreted morphogens, such as SHH, WNT, BMP, and FGF. These morphogens create spatiotemporal gene regulation patterns, leading to the formation of two-dimensional feather primordia arrays (Desmarquet-Trin Dinh and Manceau, 2025; Jung et al., 1998; Lin et al., 2009; Riddell and Headon, 2025; Turing, 1952). The initial symmetric feather buds subsequently elongate into asymmetric structures along the direction of elongation (Chen et al., 1997; Li et al., 2017) influenced by interactions between the bud epithelium and the mesenchymal zone of polarizing activity, involving pathways like Wnt, Notch, and non-muscle myosin (Li et al., 2013).

The Turing reaction-diffusion mechanism, involving the slow diffusion of morphogens over short distances may interact with high-speed electrical signaling via ion channels to influence tissue development (Daane et al., 2021; Lanni et al., 2019; Zhang and Levin, 2025). For instance, suppressing the gap junction channel, connexin 30, with an siRNA or with a dominant negative form at HH26 inhibited dermal condensation formation (Tseng et al., 2024) whereas, blocking gap junctions at HH34 induced new feather bud waves, suggesting that an inhibitor travels long distances through gap junctions at this stage (Tseng et al., 2024).Additionally, applying a stable or pulsed electric current across the chicken skin chicken skin collectively disrupts feather bud orientations, suggesting endogenous bioelectrical activity involvement in this process (Jiang et al., 2021). The genetically encoded calcium indicator GCaMP6 showed oscillating calcium activity at HH31 in short feather buds which synchronizes into travelling waves propagating from the tip of elongating HH 32-36 feather buds in a Shh/Wnt/gap junction-dependent manner. Inhibiting specific voltage-gated calcium channels, iCRAC (calcium release-activated calcium channels) or gap junction channels led to diminished calcium signaling, resulting in stunted feather development (Li et al., 2018).

We applied vibrating probe technology to developing feather buds to map extracellular electrical currents in 3 dimensions (Li et al., 2018). As diagrammed, at HH31 all currents at the dorsal skin surface flow inward (Fig. 1A, right panels). However, by HH35, as feather buds formed and elongated, an outward current emerges from the anterior region of the feather bud, while an inward current persists at the bud tip. Extracellular currents flowing adjacent to the tissue to complete a standing current loop. Calcium channel blockers demonstrated their involvement in the inward bioelectric current (Li et al., 2018), but since these channels do not facilitate outward currents we hypothesize that potassium channels mediate the outward current to complete the loop. Indeed, our previous bulk RNA-seq analyses (Fig. 1B) identified an array of potassium channels in feather bud mesenchymal cells (Li et al., 2018) and in situ RNA hybridization confirmed their localization within the anterior feather bud base (Li et al., 2018).

**Figure 1.**
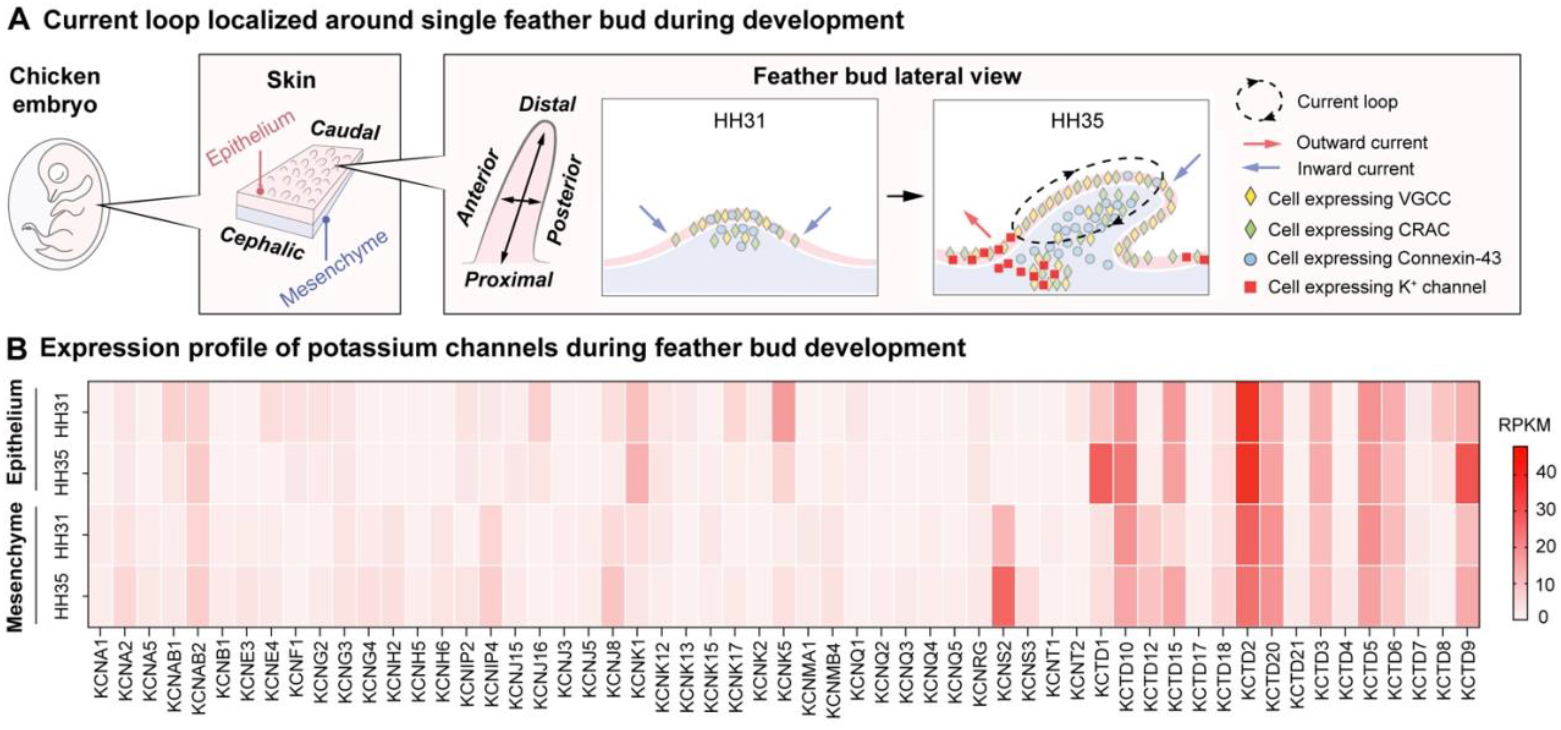
A. Diagrams of chicken embryo skin explant with defined anatomical terms. Single feather bud at HH31 and 35 (right two panels), depicting ion channels contributing to ion current and arrows indicating direction of electrical current flow (blue, inward; red outward) VGCC voltage-gated calcium channel; CRAC calcium release-activated calcium channel.Dashed arrows indicate standing current loop. B. Bulk RNAseq at HH31 and HH35. Chicken-skin epithelium and mesenchyme show K channel expression changes during development.

The mesenchymal cells in feather buds create a gap junction-coupled cell collective, allowing the fluorescent dye lucifer yellow to move between cells, validating their connectivity at HH35 (Li et al., 2018; Tseng et al., 2024). Consequently, the loop current may enter cells at the feather bud tip, travel through gap junction-coupled cells along the bud axis, exit at the anterior base, and subsequently return through the extracellular medium to the tip.

This study investigates the function of potassium channels in establishing the standing current loop in feather buds. The current loop requires diverse channel types positioned at widely separated locations to maintain the current’s flow. Blocking potassium channels should block or redirect the normal current. We also evaluate whether disrupting this electric current loop perturbs feather bud periodic patterning, elongation, and expression of molecular markers.

## Results and Discussion

### Potassium channel blockers perturb feather periodic patterning and bud elongation

We used chicken dorsal-skin explants at HH 31, 33, 35 to evaluate the role of potassium channels in feather-bud development (Fig. 2A). As there are many types of potassium channels in the developing chicken skin (Fig. 1B), we applied nonspecific K channel blockers tetraethylammonium TEA and 4-aminopyridine 4-AP (Hille, 2001).

**Figure 2.**
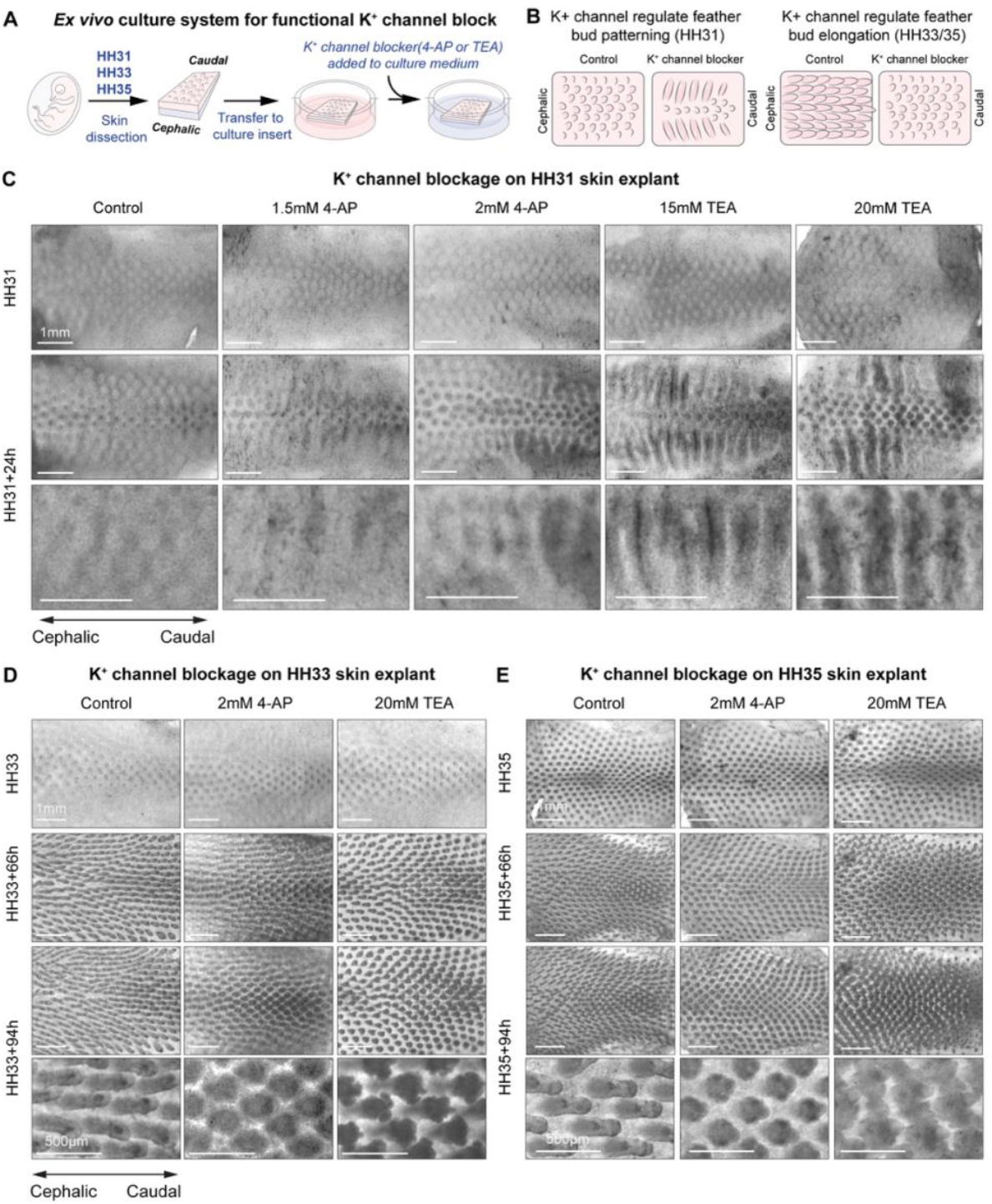
A, B. Diagram showing experimental protocol. C. Stage-matched E7/HH31 explants treated with either TEA (15 or 20 mM) or 4-AP (1.5 or 2 mM) display altered development of new, distinct buds at 24 hours of treatment, sometimes resulting in a medial-lateral stripe-like dermal condensation. Established bud development was also inhibited as center buds do not enlarge or gain height. Longer incubation (62, 86 and 110 hours) of HH31 explants with either TEA or 4-AP inhibits feather bud elongation (Supplementary data Figs. 1 and 2). D. Stage-matched E8/HH33 explants treated with either 2 mM 4-AP or 20 mM TEA for 66 or 94 hours show distinct morphological differences compared to control samples. At this stage, central buds from 4-AP and TEA treated explants were significantly shorter (**p<1.2e-08) with significantly smaller length to width aspect ratios at 66 hours (*p<0.01), and 94 hours (**p<1.2e-08), (Supplementary data Fig. 3; N=4 samples, N=7 central buds each) and were misshapen with small projections. E. Stage-matched HH35 explants treated with either 2 mM 4-AP or 20 mM TEA also show distinct morphological differences. Central buds from 4-AP and TEA treated HH35 explants were not significantly shorter at 66 hours but did show a significantly smaller aspect ratio with either 4-AP or TEA at 66 hours (*p< 3e-10). Longer treatment of 94 hours showed significantly shorter central buds with smaller aspect ratios (*p< 3e-10) (Supplementary data Fig. 4) that were also misshapen.

Applying potassium-channel blockers leads to clear dose- and developmental stage-dependent changes in bud morphogenesis. At HH31, seven to nine parallel anterior-to-posterior rows of early buds have developed. Further bud development was altered in the next 24 hours in a dose-dependent manner with 4-AP or TEA (Fig. 2). High doses of 4-AP (1.5 mM N=6, 2 mM N=29) and TEA (15mM N=6, 20 mM N=30) delayed the formation of additional rows of early buds compared to control (N =30) (Fig. 2C), and those buds merged together to form medial-lateral “stripes” (see higher magnification images in third row), whereas lower doses of 4-AP (0.5 mM N=4, 1 mM N=6) and TEA (5 mM N=4, 10 mM N=6) had little or no effect on new bud formation (data not shown). Furthermore, established buds did not grow as much as they did in the control. Photographs and quantification of bud lengths and aspect ratios after treatment at HH31 + 62, 86 and 110 hours are shown (Fig. S1 and S2).

Treating skin explants from later-stage (HH 33 and 35) chicken embryos with either 2mM 4-AP or 20 mM TEA did not show significant differences with short incubations (24 hours, not shown) but clear morphological differences were observed at 66 and 94 hours of treatment, when compared to the control. Explants from HH33 chicken embryos treated with either 4-AP (N=19) or TEA (N=18) displayed significantly shorter central buds (p<1.2e-08) with smaller aspect ratios at both 66 (p<0.01) and 94 hours (p<1.2e-08) of treatment compared to control (N=16) (Supplementary Fig. 3). Explants from later-stage HH35 chicken embryos treated with either 4-AP (N=9) or TEA (N=9) did not show a significant difference in central bud length at 66 hours but showed a decreased aspect ratio (p<3e-10) compared to control (N=8) (Supplementary Fig. 4). At 94 hours, center buds from HH35 skin explants treated with either 4-AP or TEA were significantly shorter in length than controls and displayed significantly smaller aspect ratios (p<3e-10). Furthermore, at both 66 and 94 hours and for both 4-AP and TEA, buds were not only shorter, but also misshapen, with multiple short projections, especially at high TEA levels (See enlarged images in 2D and E, row 4).

To study molecular expression in these perturbed buds, we used whole-mount in-situ RNA hybridization to analyze effects of TEA and 4-AP on the expression of beta-catenin which is enriched in the bud primordia, and SHH, which is expressed in the distal-posterior portion of the feather-bud epithelium (Noramly et al., 1999; Ting-Berreth and Chuong, 1996; Widelitz et al., 2000). We observed that while control samples exhibited the expected expression pattern of SHH, the samples treated with 20 mM TEA or 2 mM 4-AP showed reduced SHH expression at 24 hours of treatment and further reduced and wider spots of expression of SHH at 40 hours of treatment (Fig. 3A). Beta-catenin, a crucial component of Wnt/beta-catenin signaling that is known to be necessary for feather bud formation (Noramly et al., 1999; Widelitz et al., 2000), shows decreased expression in buds at 24 hours of treatment and very weak expression at 40 hours treatment when compared to the control (Fig. 3B).

**Figure 3.**
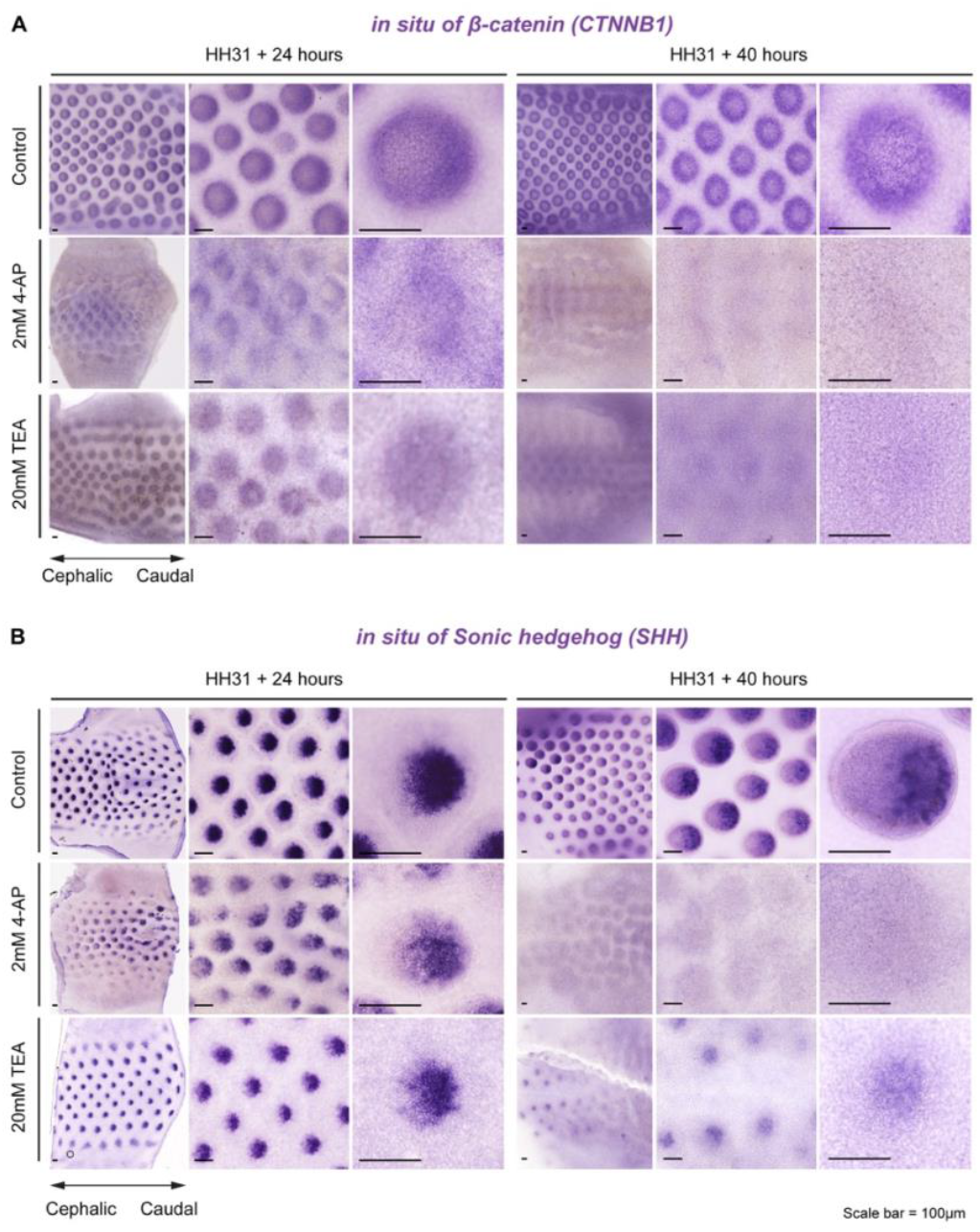
Whole-mount in situ hybridization against beta-catenin and SHH confirms that potassium channel inhibitors TEA and 4-AP inhibit initial feather-bud formation and growth. Stage-matched HH31 explants treated with either 20 mM TEA or 2 mM 4-AP display reduced SHH and beta-catenin mRNA expression at 24 hours of treatment, and expression became diffuse after 40 hours of treatment. The three images at each stage show views with progressively higher magnification.

These results show that blocking potassium channel activities reduces the expression of Sonic Hedgehog and beta-catenin, both markers for successful feather buds. The results indicate the importance of potassium-channel activity in feather bud formation.

### Potassium channel blocker TEA blocks outward current in the anterior feather bud and inward current in the posterior bud

The non-invasive vibrating probe technique was used to measure the current density and direction of standing extracellular currents around developing feather buds for HH 31-37. In Figure 4A, right side, negative values indicate current flowing into the feather bud, and positive values indicate current flowing out of the bud. Currents at the anterior (blue), and posterior (green) regions are displayed as the mean (N=10) with SEM. Anterior currents were determined to be significantly different from those at the posterior bud during HH 32-37, as the current flows in opposite directions at these locations (**p<1e-5, *p<0.05). In fact, the currents flowing into the bud at the tip were similar in magnitude to those flowing out at the base, especially for HH 32-37. That is, the current that flows into the bud also flows out of the bud (Supplementary Fig. 5).

**Figure 4.**
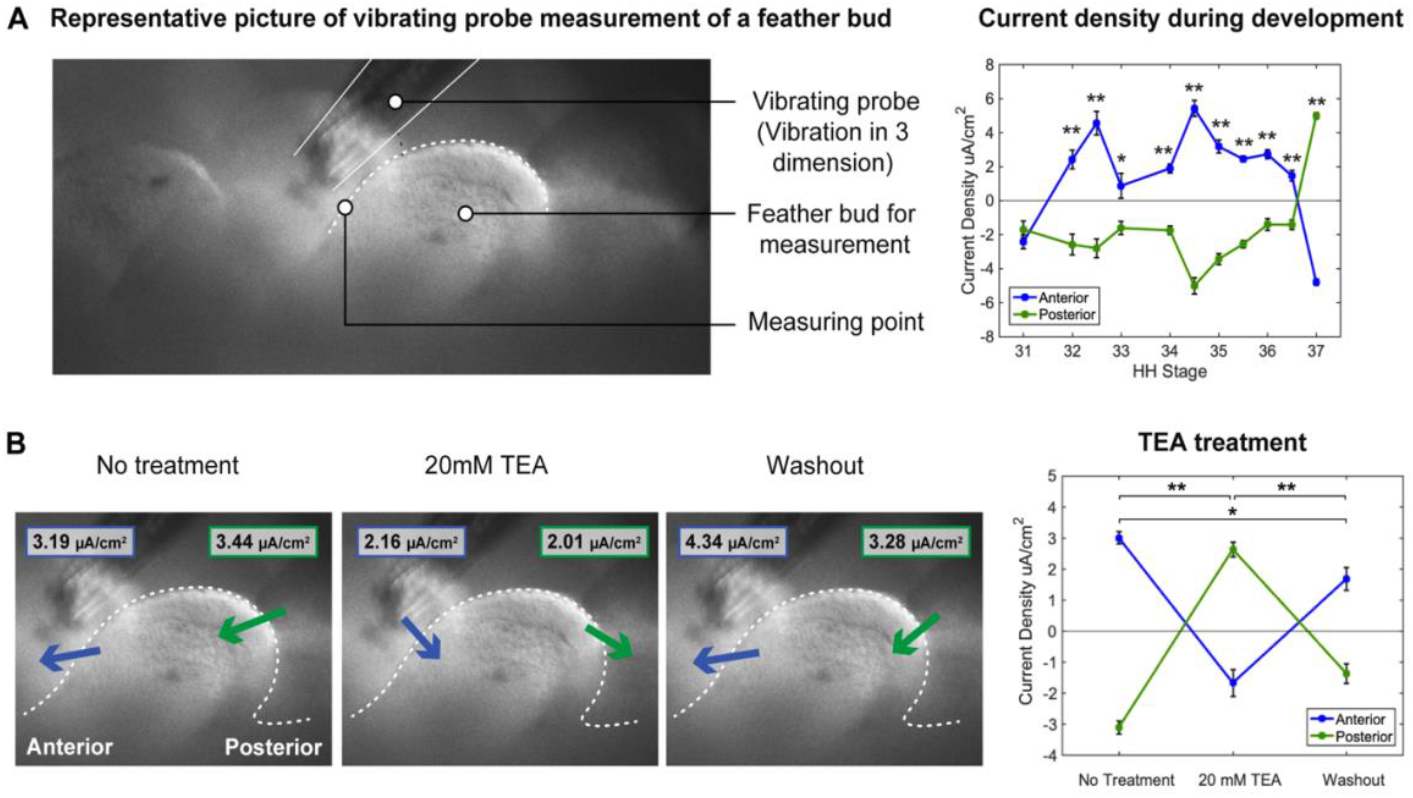
A. Image of vibrating probe (outlined by white lines) poised over the anterior end of a feather bud. The right panel summarizes current density measured at the anterior (blue) or posterior (green) bud at specified HH stages. Negative values indicate current flowing into the feather bud, and positive values indicate current flowing out of the bud. Currents are displayed as mean (N=10) and standard error of the mean (whiskers) (**p<1e-5, *p<0.05). Current is uniformly inward at HH 31. HH 32-36, current is outward at the anterior bud and inward at the posterior bud. The current “flips” direction at HH 37. B. Effect of TEA treatment on anterior and posterior current density vector. Measurements were conducted at the baseline, after 50 minutes of 20 mM TEA treatment, and after 50 minutes of TEA washout for four separate experiments (**p<1e-5, *p<0.02).

As illustrated in Fig. 4B, at HH 35-36, with 20 mM TEA treatment the outward current at the anterior feather bud base no longer flowed outward (as expected for TEA action on potassium current), and instead flowed inward. TEA also caused the inward tip current to change direction to an outward current. This is surprising, as extracellular TEA does not block sodium or calcium channels, the major ion channels responsible for inward currents. TEA action was reversible. The mutual dependency of the inward and outward currents points to their being components of the same current originally identified as a standing extracellular loop current (Li et al., 2018).

### Implications of potassium channel activity in feather morphogenesis

In summary, we demonstrate that treating HH31 chick embryo skin explants with potassium channel blockers suppresses feather bud-associated SHH and beta-catenin mRNA expression and alters feather bud patterning, converting periodic feather arrays into stripes. At HH 31-35, treatment with these blockers suppress feather bud elongation, altering the morphology of individual feather buds.

Previous research has highlighted the significance of potassium channels in embryonic tissue morphogenesis (Daane et al., 2021; George et al., 2019; Hoptak-Solga et al., 2007; Lanni et al., 2019; Perathoner et al., 2014; Podobnik et al., 2020; Stewart et al., 2021). Notably, kcnh2a regulates the length of fins in flying fish (Lanni et al., 2019) while kcnk5b serves a similar function in zebrafish (Daane et al., 2018). Additionally, inwardly rectifying potassium channels are implicated in BMP secretion essential for Drosophila wing formation (Dahal et al., 2017) and murine palate morphogenesis (Follmer et al., 2024).

The potassium channel is believed to play a crucial role in regulating the resting membrane potential and possibly calcium entry. Calcium serves dual functions as a current carrier and second messenger, participating in process such as secretion (Huizar et al., 2020), cell motility (Li et al., 2018), and gene regulation (Greer and Greenberg, 2008; West et al., 2002). The entry of calcium is voltage-dependent, occurring through voltage-gated channels and inwardly rectifying channels that primarily conduct when the membrane potential is negative (Fig. 1A).

The present study provides new insights into how potassium channels establishing a voltage gradient among cells within the gap-junction-coupled cell collective might contribute to feather bud morphogenesis by regulating electrical current flow (Fig. 1A and Fig. 4A). The membrane potential of current flowing inwards through positively charged calcium channels at the bud tip flows out the negatively charged base that expresses potassium channels (in situ RNA hybridization) (Li et al., 2018). Potassium should set the membrane potential close to −80 to −90mV, its Nernst Potential. Thus, as found with the vibrating probe, the potential gradient would be directed from the positive tip to the negative base.

TEA blocks outward potassium currents, leading the voltage at the base to become less negative. This causes the current to flow into the base and out of the feather bud tip, reversing polarity. The loss of inward current at the tip is surprising because TEA should not block inward currents. The change in the overall current direction from the base to the tip suggests that blocking potassium current causes the base to become more positive than the tip. Like a seesaw, the current direction can be determined by which end is more negative. The findings make sense when we recognize that the current at the tip and the base are components of the same macroscopic current, just measured at the points of entry and exit in the feather bud. In support of this concept, the current amplitudes measured at the tip and the base are equal in magnitude, but opposite in direction (Fig. 4A and B, Supplementary Fig. 5).

What purpose does the standing current fulfill? The standing current appears just as the feather bud begins to elongate at HH32-36. Our previous work showed that most, if not all, of the inward current at the feather bud tip is carried by voltage-gated and calcium-release activated calcium channels, which initiate calcium waves that spread rapidly and deeply into the feather bud from the tip to the base. This transport of calcium waves through gap-junction-coupled cell collective is similar to that seen in cardiac muscle, a functional syncytium.

Experimental disruption of potassium channels at the base of the feather bud flips the standing current’s direction and leads to stunted and misshapen feather buds. This deformity may stem from diminished activation of calcium-dependent atypical myosins, which enhance cell motility (Li et al, 2018), and possibly reduced morphogen secretion.

At HH37, the standing current exhibits a spontaneous reversal in direction, similar to the response seen with TEA treatment, suggesting a potential change in the voltage gradient within the bud, only this seesaw flip is spontaneous. This reversal may occur if the primary potassium channels at the base are calcium-activated, leading to channel closure due to decreased cytoplasmic calcium, thereby reducing the negativity of the base voltage.

Alternatively, the activation of new potassium or chloride channels at the feather bud tip may contribute to a more negative tip voltage. Further experimentation is necessary to elucidate the role of this current reversal, as well as to identify the specific potassium channels present at the feather bud base.

The feather bud represents a unique functional syncytium characterized by a collection of cells exhibiting regional specialization rather than uniformity. Previous in situ hybridization studies revealed stage- and region-specific expression of ion channels, such as an accumulation of calcium entry pathway transcripts at the feather bud tip, widespread dermal and epidermal gap junction expression, and potassium channels located at the feather bud base. Additionally, when isolated, feather bud mesenchymal cells showed functional heterogeneity, displaying at least four distinct cytoplasmic calcium response patterns to extracellular potassium stimulation. This cellular diversity plays a crucial role in feather bud design, generating macroscopic currents, which coordinate developmental processes at appropriate locations and developmental time points.

Our study highlights the significant role of non-neural bioelectricity in development, specifically through the action of ion channels which shape the 3D architecture of early skin appendages. Diverse channels work together in a complex multiscale mechanism that orchestrates feather bud genesis and elongation. This raises the intriguing possibility that analogous mechanisms could be involved in the development of other skin and body appendages.

## Materials and Methods

### Egg Resources

Animal care and experiments were conducted according to the guidelines established by USC Institutional Animal Care and Use Committee. White leghorn chicken eggs were purchased from Charles River (pathogen free) and AA laboratory.

### Vibrating Probe

Electrical current flowing through a resistive medium creates an “ohmic” voltage drop. The vibrating probe technique non-invasively measures the small voltage difference at a fixed separation distance (on the scale of microns) in order to calculate current density, as previously described (Li et al., 2018; Shen et al., 2016). Briefly, using piezo elements, a single voltage-sensor electrode tip is “vibrated” simultaneously in three mutually-independent directions at mutually-independent frequencies. The voltage signal output is fed into three phase-sensitive lockin detectors to separate the magnitude of currents in X-, Y-, and Z-directions.

We used a custom-designed Scanning Vibrating Electrode Technique (SVET) system (Science Wares). Under the control of custom software (Science Wares) the voltage-sensor probe is maneuvered near the tissue, using computer-controlled stepper motor stages.

Reference measurements are taken far from the tissue sample before and after recordings. The custom software allows programmed and repeatable measurements at mapped locations.

The voltage sensor is a platinum-iridium probe (Microprobes for Life Science PI10036.0A10) plated with platinum black in three steps: 2-3 minutes at −200 nA, 3-5 minutes at −500 nA until reaching 80% of desired diameter, and 3-5 intervals of 1-2 seconds at −1 uA. The higher current bursts at the end create a high-capacitance platinum-black layer on the probe. The freshly plated probe is calibrated with a set current density of 20 µA/cm^2 before and after each experiment.

Explants are mounted in custom vibrating probe dishes by cutting the explant-attached membrane from the cell culture insert using 1-mm-thick double-sided tape. Measurements on chicken-skin explants from different staged embryos are performed at room temperature in external solution containing (mM): 140 NaCl, 2.8 KCl, 2 CaCl2, 1 MgCl2, 10 HEPES, and 10 D-Glucose, pH adjusted to 7.3 with NaOH and 330 mOsm. Substitute 20 mM NaCl with equimolar TEA-Cl for potassium channel inhibition experiments.

### Chicken Skin Explant Culture

Dorsal skins from different stages of chicken embryo were carefully dissected in Hank’s Buffered Salt Solution (HBSS, Gibco) and transferred to culture inserts (Falcon, 0.4 µm pore) for imaging and vibrating-probe experiments. Tissue was grown on inserts for a minimum of four hours before use. Culture Medium consisted of high-glucose DMEM with Glutamax supplemented with 10% FBS, 2% Chicken Serum, and 1% Pen/Strep (all Gibco). For potassium-channel inhibition, media was supplemented with either 5-20 mM Tetraethylammonium chloride (TEA, Sigma) or 0.5-2 mM 4-Aminopyridine (4-AP, Sigma).

### Confocal Imaging

Confocal imaging of RCAS-GCaMP6s-2A-mCherry of E9/HH35 was performed on a Leica Stellaris-5 Confocal at 37°C/65%Humidity/5% CO2 (oko-lab). Imaging media consisted of FluoroBright high-glucose DMEM supplemented with Glutamax, 10% FBS, 2% Chicken Serum, and 1% Pen/Strep (all Gibco). For potassium-channel inhibition, media was supplemented with 20 mM TEA (Sigma). ImageJ was used for imaging analysis. ROIs (N>20 for each sample) were selected within a bud and fluorescence intensity was recorded with no treatment for 25 minutes, followed by TEA treatment for 30 minutes before recording for another 25 minutes. Matlab was used to determine peaks in Calcium signaling using a MinPeakHeight threshold of 20. Data is displayed as a boxplot

### Virus Production and Injection

Virus production of RCAS-GCaMP6s-2A-mCherry has been reported (Li et al., 2018). Virus was concentrated by ultra-centrifuging in a BECKMAN Coulter L8–80M (SW28 rotor) at 26000 rpm for 1.5 h. Concentrated virus was then injected into H&H 14 chicken embryo somites for dermal expression.

### In situ hybridization

In situ hybridization against Sonic Hedgehog (SHH) and beta-catenin was performed as published previously (Chuong et al., 1996; Widelitz et al., 2000). Primers used to produce the probes have also been reported previously (Li et al., 2018).

### Statistical Analysis

Independent Student’s T-tests were performed on all data, except where noted. In those cases, Paired Student’s T-tests were performed. Significance represented by p value. Only a p<0.05 was considered to be statistically significant.

### Single-cell Mesenchymal Culture

Dorsal Skins from E7-E9 were incubated in cold 2X Calcium-Magnesium Free PBS (CMF) supplemented with 0.25% EDTA for ten to twenty minutes. The epithelial layer was manually removed using fine forceps. The remaining mesenchymal tissue was rinsed in HBSS before dissociation in 0.35% type 1 Collagenase (Worthington) for twenty minutes at 37°C. The digestion was halted by adding equal volume of culture media (same as explant). Cells were reconstituted in culture media and plated on custom glass-bottom dishes coated with 10% Type 1 Collagen (Sigma) in plain DMEM for 30 minutes at room temperature. Cells were cultured 8-24 hours at 37°C/5% CO2 before performing imaging or patch-clamp experiments.

### RNAseq

Skins dissected from H&H 31 and H&H 35 chicken embryos were used for RNA-Seq library preparation. Epithelial and mesenchymal tissues were separated in 2X CMF (calcium-magnesium free saline). Total RNA was extracted using Trizol reagent. For library construction we used TruSeq RNA Sample Prep Kit Version2 (Illumina). Two biological replicates were sent for sequencing (50 bp, single end) with Illumina HiSeq 2000 in the USC Epigenome Center. FastQ files were trimmed and mapped to the chicken genome (galGal4) using Partek Flow. Further analysis was done in Partek Genomic Suite.

## Acknowledgements

We thank Eric Karplus at Sciencewares.com for setting up the vibrating probe apparatus for our specific needs.

