## Supplementary Information for "Roles of K-channel activity in feather bud morphogenesis"

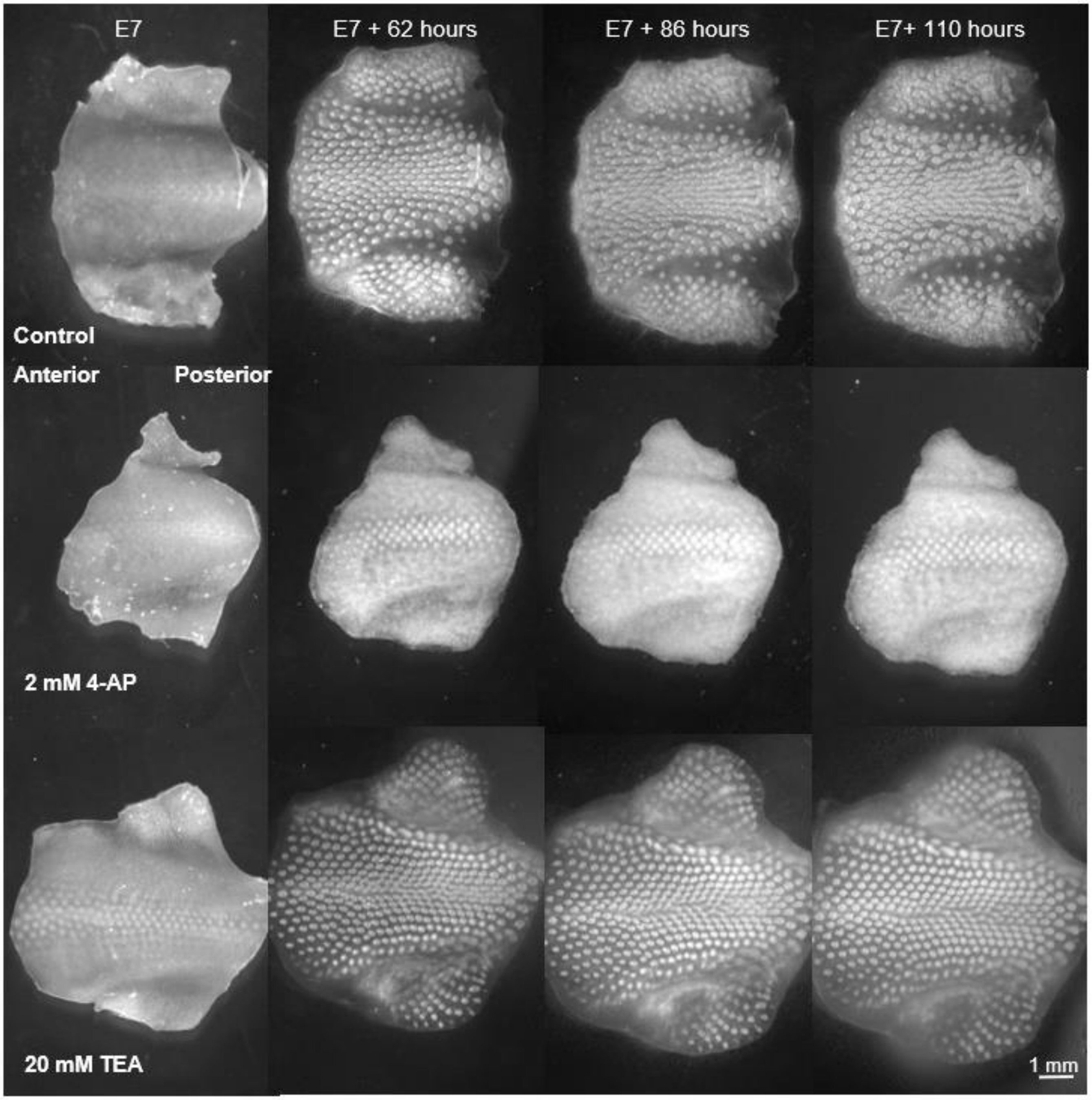


**Figure S1. Brightfield images of E7/HH31 chick skin explants, control and 2 mM 4-AP or 20 mM TEA treatments at 0, 62, 86, and 110 hours.**

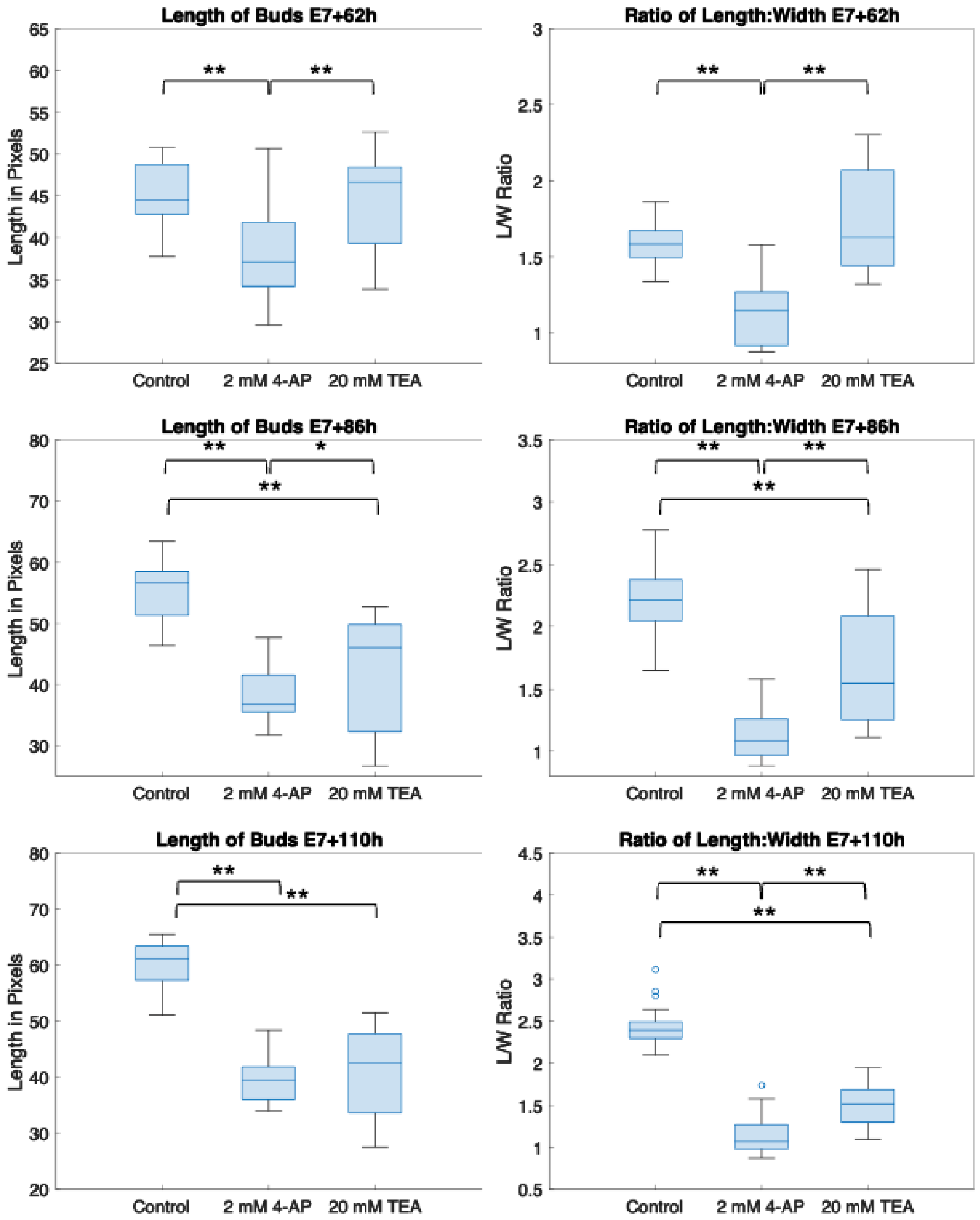


**Figure S2. Potassium channel inhibitors 4-AP and TEA inhibit feather-bud elongation at HH31.**

**A.** Stage-matched HH31 explants treated with either 2 mM 4-AP or 20 mM TEA treated for 62, 86, and 110 hours show distinct morphological differences such as the lack of bud lengthening with TEA treatment, and the complete lack of further bud development as seen with the 4-AP treatment **B.** Central buds from TEA and 4-AP treatment were significantly shorter with significantly smaller length to width aspect ratios (*p<0.0005 **p<0.05), (N=4 samples per condition, n=7 central buds each sample)

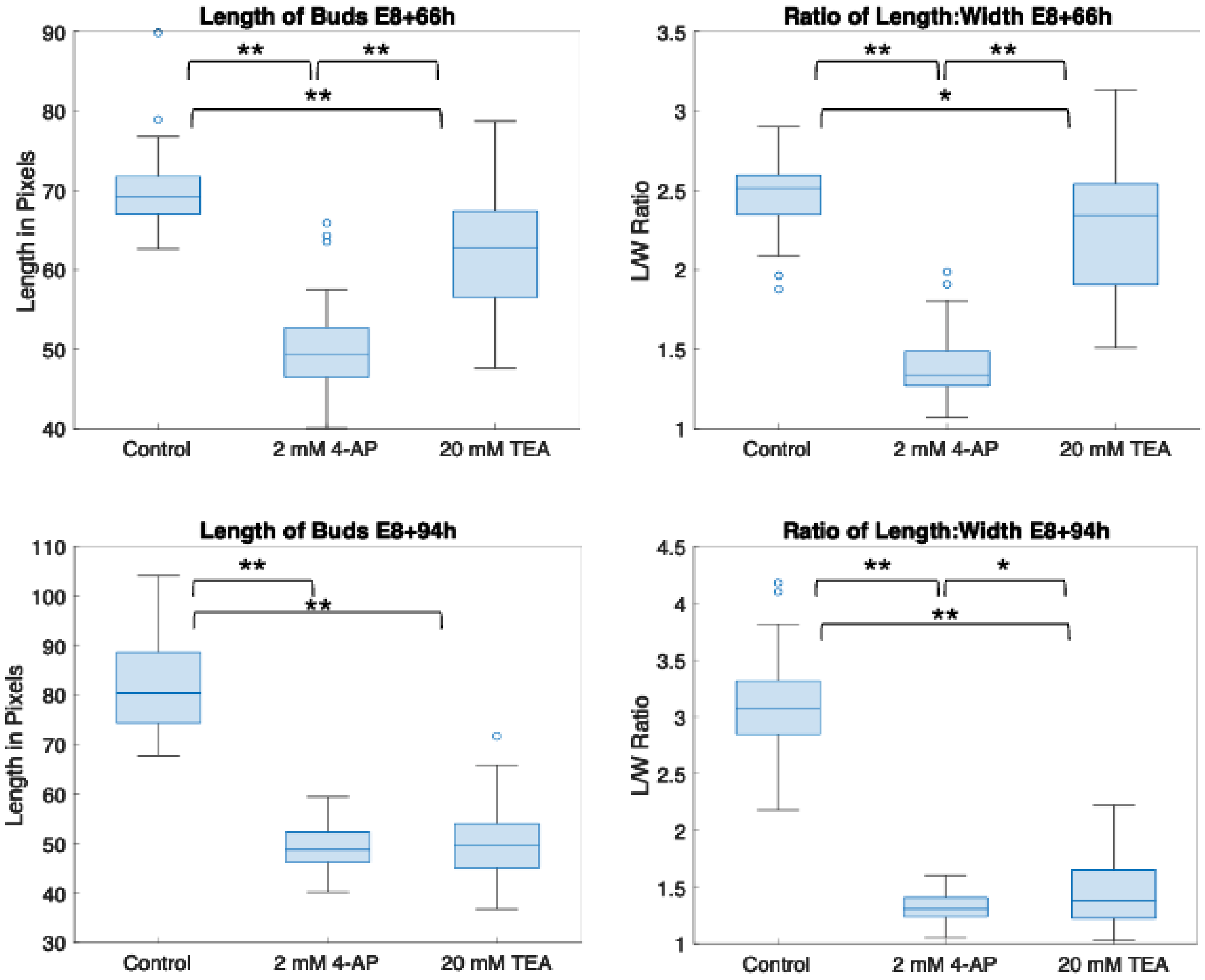


**Figure S3. Potassium channel inhibitors 4-AP and TEA inhibit feather-bud elongation at E8/HH33.** Stage-matched (E8/HH33) explants treated with either 2 mM 4-AP or 20 mM TEA for 66 or 94 hours show significantly shorter central buds (**p<1.2e-08) with significantly smaller length to width aspect ratios at 66 hours (*p<0.01), and at 94 hours (**p<1.2e-08) (N=3 samples per condition, n=14 central buds each sample)

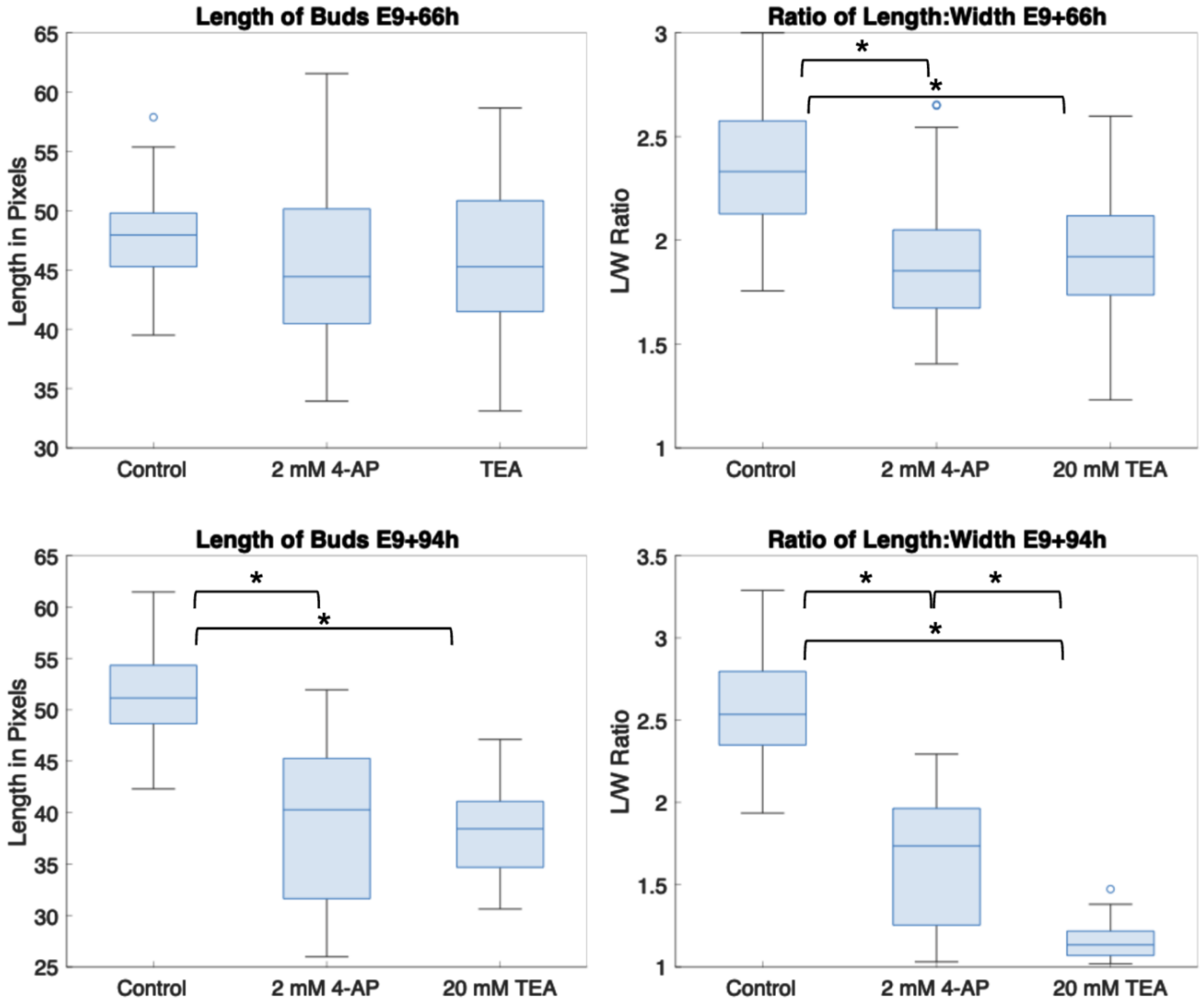


**Figure S4. Potassium channel inhibitors 4-AP and TEA inhibit feather-bud elongation at E9/HH35.** Stage-matched (E9/HH35) explants treated with either 2 mM 4-AP or 20 mM TEA for 66 or 94 hours also show distinct morphological differences. Central buds from 4-AP and TEA treated E9/HH35 explants were not significantly shorter at 66 hours but did show a significantly smaller aspect ratio with either 4-AP or TEA at 66 hours (*p< 3e-10). Longer treatment of 94 hours also showed significantly shorter central buds with smaller aspect ratios (*p< 3e-10) (N=3 samples each condition, n=14 central buds each sample).

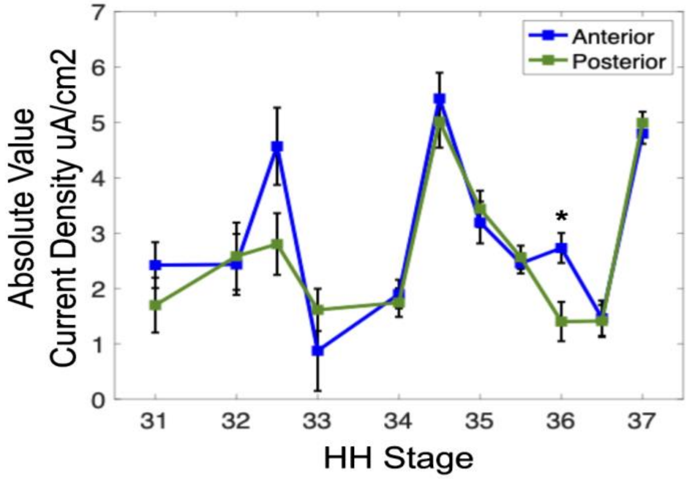


**Figure S5. Magnitude of the current density measured at bud anterior (blue) or posterior (green) at specified HH stages.** Currents are displayed as mean (N=10) and standard error of the mean (whiskers). Only at HH 36 are the currents statistically different (student's t test $p<0.05$).
